# Multifaceted regulations of HSV-1 ICP0 on the Anti-Viral Restrictions Imposed by the Host Hippo Kinases Reprogramming

**DOI:** 10.64898/2026.06.11.731592

**Authors:** Brendyn M. St. Louis, Udayan Guha, Pei-Chung Lee, Haidong Gu

## Abstract

The Hippo pathway is conserved across eukaryotes and controls key biological processes, including cell growth and organ development. Notably, humans with biallelic loss-of-function mutations in the Hippo kinase gene *MST1* suffer from combined immunodeficiency, including recurrent infections of herpes simplex virus (HSV), implicating the pathway in host immune regulation. We investigated the role of MST1 and its homolog MST2 in HSV-1 infection. We found that human epithelial cells HEp-2 proteolytically converted full-length MST1/2 into smaller N-terminal fragments (MST1/2-NT) to enhance cell apoptosis in response to the HSV-1 infection. Moreover, while infection by mutants lacking ICP0 or US3 elevated the production of MST1/2-NT and apoptosis, the overexpression of MST1-NT significantly reduced HSV-1 replication, revealing anti-viral properties of MST1/2 cleavage and the viral counteractions by ICP0 and US3. Consistently, we discovered that MST1/2-NT production, which was high in ΔICP0-infected HEp-2 cells, was completely diminished in cells permissive to the ΔICP0 infection, linking the counteraction against host MST1/2 cleavage to ICP0 functions. In addition, host caspases cleaved MST1/2 with differential preferences toward MST1 or MST2 in different infection contexts, indicating multiple regulations on the MST1/2-NT production during HSV-1 infection. While double-knockouts of MST1/2 in HEp-2 cells had marginal effects on wild type HSV-1 replication, it substantially reduced the early intake of ΔICP0 DNA, suggesting a role of full-length MST1/2 in early infection. Altogether, these results uncover the novel and distinct roles of full-length and cleaved Hippo kinases during HSV-1 infection and their multifaceted interactions with ICP0.

**Importance:** HSV poses serious threats to human health, ranging from cold sores to fatal brain infection. As a successful human pathogen, it deploys various viral proteins to counteract host defenses and subjugate host machinery, but mechanisms underlying these complex HSV-host interactions are not fully understood. For the first time, we report multifaceted interactions between the Hippo kinases and viral proteins during HSV-1 infection. We show that the Hippo kinases are converted to smaller fragments through protein cleavage in HSV-1 infected cells, and the cleaved Hippo fragments are accompanied by host cell death to execute their anti-viral activities. Moreover, multiple viral proteins contribute to counteracting this host defense, including ICP0, which executes a complex interplay with the Hippo kinases to promote infection. Understanding this new layer of virus-host interaction may pave the road to developing novel treatments for herpetic diseases.

## Introduction

The Hippo signaling pathway is evolutionarily conserved in eukaryotic organisms and responds to a diverse array of environmental cues, including biochemical stimuli and mechanical signals. Mammalian sterile 20-like kinases 1 (MST1) and its homolog MST2 are the core components of Hippo pathway and are well-known for their activities in limiting organ size and tumor formation in mammals. To activate the Hippo pathway, MST1/2 phosphorylate the scaffold protein, Mps one binder kinase activator-like 1 (MOB1), and the large tumor suppressor kinases (LATS), allowing formation of a MOB1-LATS complex. The MOB1-LATS1 kinase complex then phosphorylates the transcription regulator Yes-associated protein (YAP) and transcriptional co-activator with PDZ-binding motif (TAZ), which consequently alters the expression of genes important for cell growth and differentiation (1, 2).

Beyond activating the canonical MOB1/LATS/YAP signaling cascade for organ development and tumor suppression, emerging evidence has implicated the importance of MST1/2 in immune responses (3). Clinical case studies have shown that humans carrying biallelic loss-of-function mutations in *MST1* suffer from combined immune deficiency and various recurrent infections (4–6). Mice with conditional deletions of *Mst1* and *Mst2* also show abnormal immune cell development and functions (7, 8). Recent studies have revealed broad but complex implications of the Hippo components in viral pathogenesis. For example, infection of the carcinogenic human papillomavirus (HPV) strain 18 is associated with YAP/TAZ upregulation, a condition similar to Hippo “off”, in human biopsies (9). Consistently, TAZ knockdown, which mimics the Hippo “on” status, reduces the proliferation of host cells infected by HPV or Epstein-Barr virus (EBV) (9, 10), suggesting that carcinogenesis resulted from DNA virus infection is in line with the regulatory activity of canonical Hippo signaling in cancers. However, investigations in viral replication have shown diverse effects of the Hippo components on the infection phases of different viruses. Overexpression of YAP significantly reduces the nuclear entry of human cytomegalovirus (HCMV) in early infection, leading to impaired HCMV production (11). However, in stem cell-derived pancreas β-cells, the enhanced YAP production promotes the replication of coxsackievirus B (CVB) (12). Moreover, while phosphorylation of YAP/TAZ in the canonical Hippo “on” inhibits the egress of Ebola virus (13), SARS-CoV2 infection causes the Hippo “on” status but inhibition of MST1/2 or knocking down LATS promotes SARS-CoV2 replication (14). These seemingly conflicting observations imply that the canonical Hippo pathway regulates viral infections in virus and host-dependent fashions, of which the mechanisms remain poorly defined.

Herpes simplex virus 1 (HSV-1) is a ubiquitous DNA virus that infects 70% of the adult population. Upon primary infection of mucosal epithelium, HSV-1 is transported to trigeminal ganglia to establish a lifelong latency. Through sporadic reactivations, the virus achieves a massive spread and causes a wide array of human diseases, ranging from self-limited skin lesions to deadly encephalitis (15). Notably, recurrent HSV-1 infection is one of the major clinical manifestations in MST1-deficient patients (4), indicating a role of this conserved kinase in host defense against HSV-1 that has not been investigated.

Recently, a non-canonical Hippo reprogramming process, in which MST1/2 undergo proteolytic cleavage to promote programmed cell death in immune cells has been reported (16, 17). Upon challenge of the intracellular bacterium *Legionella pneumophila*, macrophages cleave MST1/2, producing ∼33 kDa N-terminal fragments (MST1/2-NT) that contain the active kinase domains. This cleavage is catalyzed by caspase-1, a cysteine protease activated by the immune sensors and inflammasomes that detect the molecular patterns from pathogens or damaged host cells. By producing MST1/2-NT, molecular death processes associated with apoptosis are enhanced, including caspase-3 activation, PARP1 cleavage, and histone H2AX phosphorylation. Importantly, MST1/2 double-knockout (DKO) macrophages have dysregulated cytokine release and become defective in triggering apoptosis to restrict the intracellular replication of *L. pneumophila* (16, 17).

In this study, we examined the dynamics of MST1/2 during HSV-1 infection. We discovered that human epithelial cells reprogram MST1/2 into the pro-apoptotic MST1/2-NT in response to HSV-1 infection similar to that in macrophages infected by *L. pneumophila*. As a counter measurement, viral proteins, including ICP0 and US3, possess novel activities that suppress the cleavage of MST1/2 in host cells. Intriguingly, MST1/2 knockout cells reveal a previously unknown effect of full-length MST1/2 on early HSV-1 infection. The results illustrate multi-level interactions between the host Hippo kinases and HSV-1 that involves the full-length MST1/2, the reprogrammed MST1/2-NT, and the viral protein ICP0 and US3.

## Results

### Human epithelial cells reprogram the Hippo kinases upon challenge of HSV-1

To determine the dynamics of host Hippo kinases during HSV-1 infection, we infected human epithelial HEp-2 cells with the prototype HSV-1(F) strain. In mock infected cells, trace amounts of MST1/2-NT and the apoptotic marker PARP1 p89 were detected after 24 hours, indicating spontaneous cleavage and apoptosis likely due to cell confluence (Figure 1A, lane 3). Interestingly, MST1/2-NT production was mildly increased upon HSV-1(F) infection at 24 hours post infection (hpi) but not at earlier time points. MST1/2-NT accumulation was associated with increasing the multiplicity of infection (MOI) from 0.1 to 1 pfu/cell but saturated at 10 pfu/cell, suggesting that the abundance of viral protein(s) overcomes the cleavage at high MOIs. Notably, the extent of PARP1 cleavage, a hallmark of apoptotic cell death, was well aligned with the amount of MST1/2-NT in the infected cells at different MOIs and timepoints (Figure 1A), which is similar to the observations in mouse macrophages undergoing apoptosis in response to chemical stimuli or the intracellular bacterium *L. pneumophila* (16, 17). These results from the non-immune cell line HEp-2 show that human epithelial cells are capable of reprogramming the Hippo kinases in response to HSV-1 infection and a viral counteraction(s) diminishes this host response at a higher dosage.

**Figure 1.**
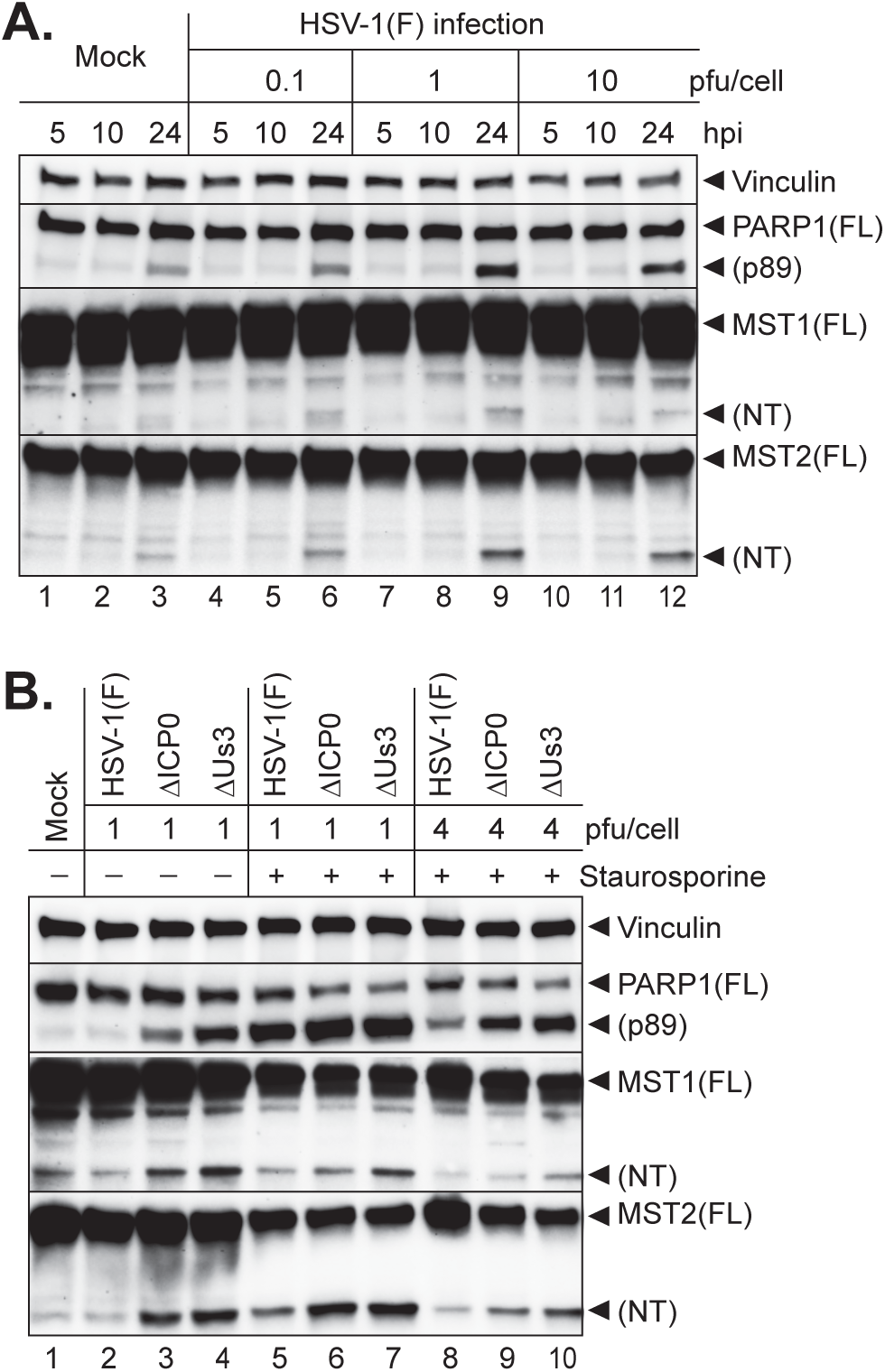
HSV-1 infection triggers MST1/2 cleavage and apoptosis. HEp-2 cells were infected with HSV-1(F) at 0.1,1 and 10 pful/cell (A) or treated with DMSO or 0.5 μM of Staurosporine for 30 minutes before infection with HSV-1(F), ΔICP0, or ΔUS3 mutant at indicated pfu/cell (B) for 24 hours. Total cell lysates were electrophoretically separated and immunoblotted for the indicated proteins.

### ICP0 and US3 impede MST1/2 reprogramming during HSV-1 infection

Programmed cell death is a critical host intrinsic defense mechanism because it deprives the intracellular pathogens of the cellular machineries essential for their growth. To ensure a successful infection, HSV-1 devotes multiple viral proteins to suppress various forms of programmed cell death, including apoptosis, necroptosis, and pyroptosis (18–22). Among them, the viral kinase US3 is well characterized for its anti-apoptotic activity by phosphorylating Bcl2 associated agonist of cell death (BAD) to block the cleavage and activation of this pro-apoptotic protein (18, 19). Another example is the viral E3 ubiquitin ligase ICP0, which inhibits the cell pyroptosis triggered by the NLRP1 inflammasome (22). Since the production of MST1/2-NT influences apoptosis and pyroptosis in macrophages challenged with the bacterial pathogen *L. pneumophila* (17), we investigated the role of US3 and ICP0 in MST1/2 cleavage by infecting HEp-2 cells with a US3-null virus (ΔUS3) or ICP0-null virus (ΔICP0). Remarkably, the deletion of either ICP0 or US3 substantially increased the production of MST1/2-NT and PARP1 p89 in cells infected at 1 pfu/cell for 24 hours (Figure 1B, lanes 2-4), suggesting that ICP0 and US3 of HSV-1 possess activities to block MST1/2 cleavage and host apoptosis.

Staurosporine is a potent apoptosis inducer which broadly inhibits protein kinases to disrupt cellular signaling (23). To characterize the relation between apoptosis and MST1/2 in HSV-1 infection, we examined the cleavage events for MST1/2 and PARP1 in staurosporine-treated HEp-2 cells. As expected, the staurosporine treatment enhanced PARP1 cleavage in cells infected with 1 pfu/cell of HSV-1(F), but this enhancement was largely suppressed by the F virus at 4 pfu/cell (Figure 1B, lanes 2, 5 and 8). Similar to the patterns of PARP1 p89, staurosporine also promoted MST2 cleavage while its effect on MST1 cleavage was limited (Figure 1B, lane 2 vs. 5), indicating the two homologous Hippo kinases may be cleaved via different mechanisms. Importantly, unlike the suppression observed in HSV-1(F) infected cells, the levels of MST2-NT and PARP1 p89 remained high in cells infected with ΔICP0 or ΔUS3 virus at 4 pfu/cell (Figure 1B, lanes 8-10). These results indicate that ICP0 and US3 block MST1/2 cleavage and apoptosis in a viral dosage-dependent manner.

### Overexpression of the MST1-NT fragment significantly blocks HSV-1 replication

To evaluate whether the MST1/2 cleavage products have a role in HSV-1 replication, we constructed plasmids carrying the full-length (FL) or N-terminal residues 1-326 (NT) of human MST1 cDNA and expressed them in HEp-2 cells by transient transfection. Compared to the mock transfection without DNA, transfection of the empty vector (EV) caused a slight increase in PARP1 cleavage (Figure 2A, lanes 1,4,7 vs. 10-12), likely due to host autonomous responses to the influx of foreign DNA, whereas transfection of either MST1-FL or MST1-NT significantly enhanced the PARP1 cleavage. We noticed that the ectopic MST1-FL was extensively cleaved at high abundance and produced MST1-NT at a level similar to that directly expressed from the MST1-NT construct, leading to comparable levels of PARP1 cleavage in the condition without HSV-1 infection (Figure 2A, lanes 2 and 3), confirming the activity of MST1-NT in promoting host apoptosis.

**Figure 2.**
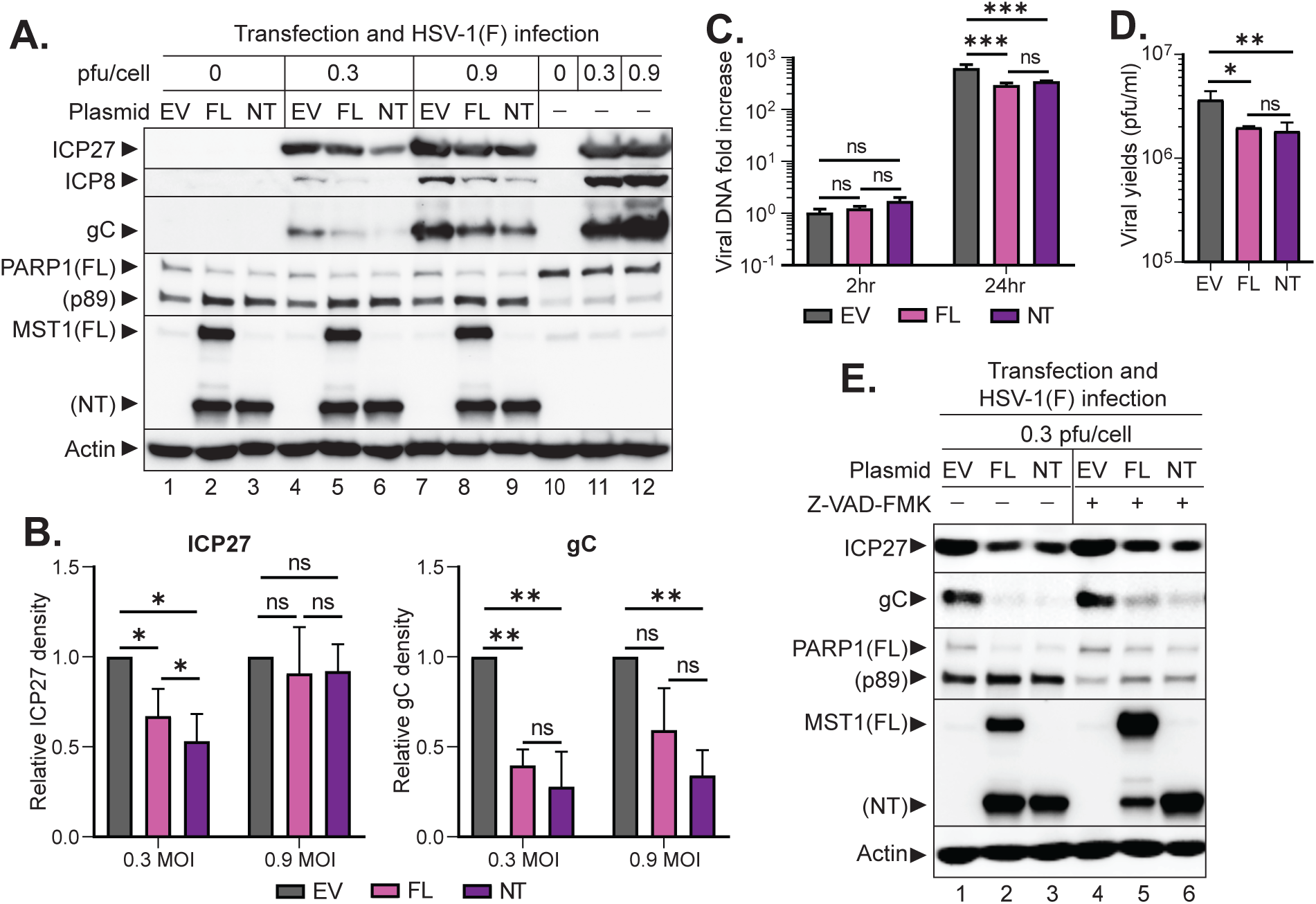
MST1-NT production inhibits HSV-1 replication. HEp-2 cells were transfected with the empty vector (EV), or plasmid expressing full-length (FL) or N-terminal fragment (NT) of MST1 for 24 hours before the following treatments. A) Transfected cells were infected by HSV-1(F) at 0.3 or 0.9 pfu/cell for 24 hours and immunoblotted for the indicated proteins. B) Band densities of ICP27 and gC in (A) were normalized to Actin. Fold changes in FL and NT samples relative to that of EV were plotted in GraphPad. C) Transfected cells were infected by HSV-1(F) at 0.3 pfu/cell. Total DNA extracted at 2 and 24 hours were subjected to comparative qPCR to measure the viral DNA fold increase relative to the EV-2 hpi sample. D) Transfected cells were infected by HSV-1(F) at 0.1 pfu/cell for 24 hours to measure the viral titer. E) Transfected cells were treated with DMSO or 100 mM of Z-VAD-FMK for 30 minutes and then infected by HSV-1(F) at 0.3 pfu/cell for 24 hours and immunoblotted for the indicated proteins. Four repeats (B,C) or three repeats (D) were analyzed by two-way ANOVA (B) or one-way ANOVA (C,D). *: P<0.05; **: P<0.01.

Next, we infected the transfected cells with HSV-1(F) at 0.3 or 0.9 pfu/cell and examined the expression of immediate early protein ICP27, early protein ICP8, and late protein gC at 24 hpi. While these viral proteins were readily detected in EV-transfected HEp-2 cells, we found that the overproduction of MST1-FL or MST1-NT severely suppressed the levels of viral proteins (Figure 2A, lanes 4-9). Quantification of the ICP27 and gC bands in the immunoblots showed significant reduction of both ICP27 and gC at 0.3 pfu/cell in cells expressing MST1-FL and MST1-NT. At 0.9 pfu/cell, ICP27 level was no longer affected by MST-FL or -NT overexpression, but the gC expression was still reduced (Figure 2B). The profound inhibition on the true late protein gC expression suggests that the inhibitory effect of MST1-NT is likely imposed on viral DNA replication, which can be saturated by increasing the viral dosage. Consistent with reduced viral protein expression, MST1-FL or -NT transfection also significantly curtailed the levels of viral DNA fold increase and virion production at 24 hpi (Figure 2C and 2D). These results suggest that the MST1 N-terminal fragment (MST1-NT) by itself has anti-viral activities against HSV-1 infection.

The abundance of MST1-NT in HEp-2 cells overexpressing full-length MST1 prompted us to investigate the influences of this cleavage process on HSV-1 infection. Treating the transfected cells with a pan-caspase inhibitor, Z-VAD-FMK, prior to infection markedly reduced the level of MST1-NT in cells transfected with the MST1-FL construct, confirming the involvement of caspases in cleaving the full-length MST1 protein. As a result from the reduction of MST1-NT, the pan-caspase inhibitor treatment improved gC expression while PARP1 cleavage was reduced (Figure 2E, lane 2 vs. 5). Collectively, these results indicate that the anti-viral activities of MST1 lie within the cleaved variant and the proteolytic process is catalyzed by host caspases.

### ICP0 by itself induces apoptosis and MST1/2 cleavage through pathways common in HEp-2 and U2OS cells

Unlike the well characterized anti-apoptotic activity of US3, the role of ICP0 in host apoptosis during HSV-1 infection is unclear. A prior study showed that a recombinant HSV-1 virus carrying ICP0 as the only immediate early protein induces host apoptosis during infection (24). In contrast, our experiment using the ΔICP0 recombinant virus showed that ICP0 is required to suppress the production of MST1/2-NT and the apoptotic marker PARP1 p89 in infected cells (Figure 1B). To further elucidate the contexts affecting the role of ICP0 in apoptosis and MST1/2 cleavage, we transiently transfected HEp-2 cells with plasmids encoding ICP0 or US3. As a control, US3 overexpression had minimal effects on apoptosis and MST1/2 cleavage (Figure 3A). We found that ICP0 overexpression induced strong cleavage of PARP1 and MST1/2, comparable to the levels triggered by Raptinal, a potent chemical activator of the intrinsic mitochondrial apoptotic pathway (25) (Figure 3A, lanes 2 and 4), indicating that ICP0 alone may disrupt cellular functions in HEp-2 cells and cause apoptosis and MST1/2 cleavage.

**Figure 3.**
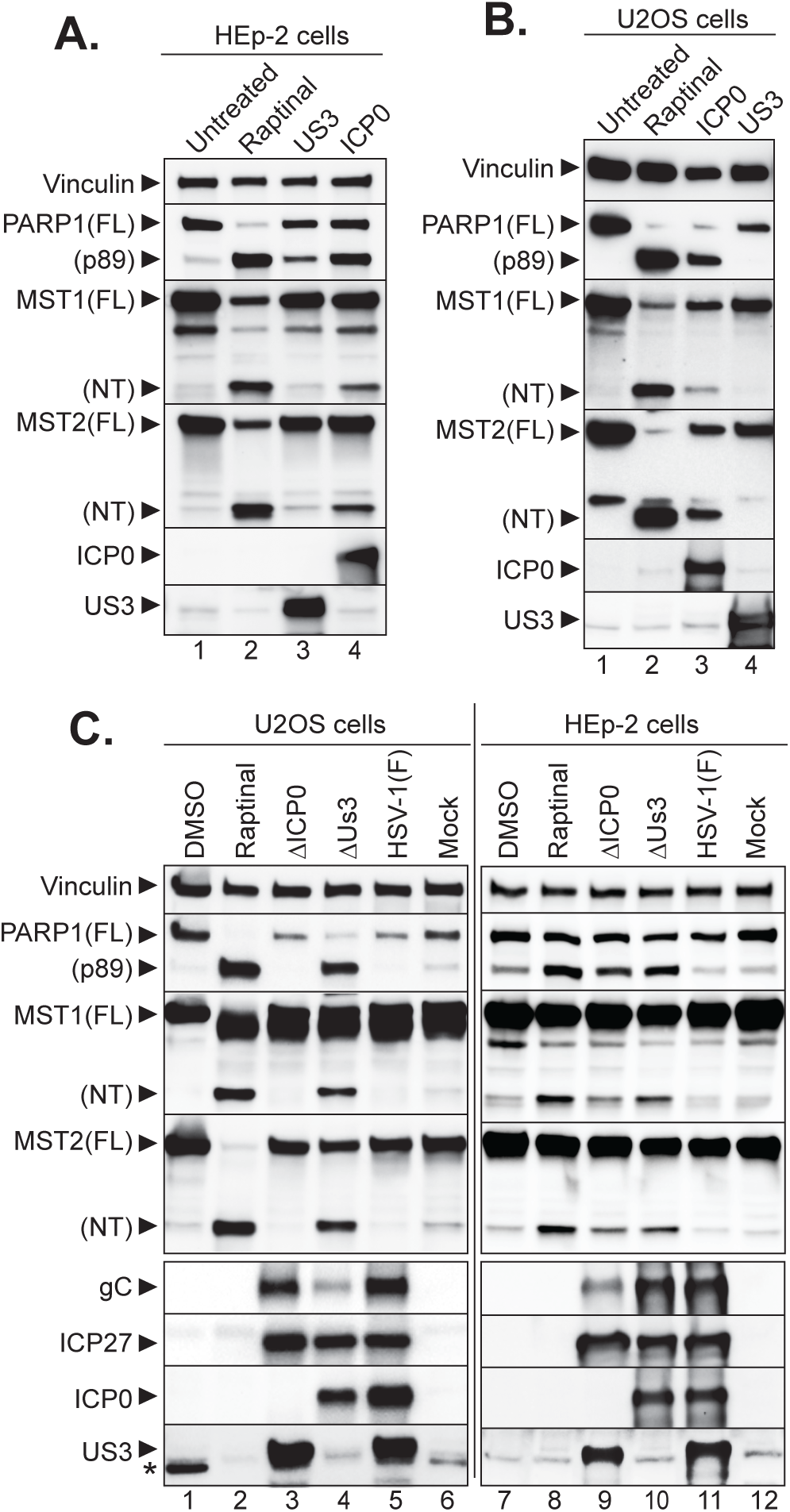
Dual actions of ICP0 in promoting or inhibiting MST1/2 cleavage associated apoptosis. HEp-2 (A) and U2OS (B) cells were treated with Raptinal or transfected with plasmids expressing ICP0 or US3 for 24 hours. C) HEp-2 and U2OS cells were mock infected, or infected by HSV-1(F), ΔICP0, or ΔUS3 mutant virus at 1 pfu/cell for 24 hours. Total cell lysates were subjected to immunoblotting for indicated cellular and viral proteins. Asterisk marks a nonspecific protein cross-reacted with the anti-US3 antibody.

ICP0 promotes the expression of viral genes by counteracting the host anti-viral restrictions imposed on the incoming DNA (26). In most cultured cells like HEp-2, HSV-1 replication is severely stalled in the absence of ICP0 at low MOI (27). U2OS, an osteosarcoma cell line defective in certain restrictive pathways, is permissive to the loss of ICP0 and allows the ΔICP0 mutant to replicate as effectively as the wild type HSV-1 at low MOI (28). To test whether U2OS cells are defective in triggering apoptosis and MST1/2 cleavage, we performed the transfection experiments in U2OS cells. Interestingly, ICP0 overexpression by itself and Raptinal treatment also induced strong cleavage of PARP1 and MST1/2 in U2OS cells while US3 overexpression had little effects (Figure 3B). These results indicate that U2OS cells are competent in activating MST1/2 cleavage and apoptosis and ICP0 alone triggers these host reactions via pathways common in both HEp-2 and U2OS cells.

### Mitigation of MST1/2 cleavage by ICP0 enhances HSV-1 replication in restrictive HEp-2 cells but not in permissive U2OS cells

Early in HSV-1 infection, ICP0 plays a critical role in antagonizing the host intrinsic/innate anti-viral defenses during HSV-1 infection (29). Two main defensive pathways targeted by ICP0 are chromatin repression of viral DNA and interferon (IFN) responses, which are missing in U2OS cells due to the lack of chromatin remodeler ATRX (Alpha-thalassemia/mental retardation syndrome X-linked) and defects in the STING induced IFN production (30, 31). The loss of these ICP0 targets renders ICP0 functions unnecessary for viral replication in U2OS cells.

Since U2OS cells express the components for executing MST1/2 cleavage and apoptosis (Figure 3B), we tested whether U2OS cells respond to the ΔICP0 virus with MST1/2 cleavage and apoptosis like HEp2 cells do (Figure 1B). Similar to the responses in HEp-2 cells, MST1/2-NT and PARP1 p89 were induced by the ΔUS3 infection but not HSV-1(F) infection in U2OS cells (Figure 3C, lanes 4-5 and 10-11), again confirming the importance of US3 in blocking apoptosis and the presence of key components for apoptosis and MST1/2 cleavage. In a stark contrast to the observations in the ΔICP0-infected HEp-2 cells, ΔICP0-infected U2OS cells did not induce MST1/2 cleavage nor apoptosis (Figure 3C, lane 3 vs. lane 9), suggesting that the defects in U2OS cells causing the permissiveness are linked to the apoptosis and MST1/2 cleavage pathway mitigated by ICP0 but not US3.

We further evaluate the impacts of MST1/2 cleavage on viral replication through the expression of viral proteins. In line with the permissiveness specific to ΔICP0 virus, we found that the lack of MST1/2 cleavage and apoptosis in ΔICP0-infected U2OS cells was correlated with a robust gC expression comparable to that in HSV-1(F) infection. Furthermore, the robust production of PARP1 p89 and MST1/2-NT in ΔUS3-infected U2OS cells was associated with a severe reduction in gC expression despite the comparable levels of immediate early protein ICP27 in all three infections (Figure 3C, lane 3-5). Contrary to U2OS cells, in the HEp-2 cells that are nonpermissive to ΔICP0 virus but permissive to ΔUS3 virus, the production of MST1/2-NT and PARP1 p89 substantially reduced gC expression in ΔICP0 infected cells but had minimal effects in the ΔUS3 infection (Figure 3C, lanes 9-10). The consistency among MST1/2-NT production, PARP1 cleavage and reduced late protein expression confirms the anti-viral effects of MST1/2 cleavage and the associated apoptosis. Furthermore, ICP0 and US3 likely block MST1/2 cleavage and apoptosis through different checkpoints, whereas the position targeted by ICP0 is linked to the missing restrictions constituting the permissiveness of U2OS cells.

### Multiple functional domains of ICP0 influence its ability to overcome the MST1/2 cleavage and apoptosis

ICP0 has multiple functional domains, including a RING-type E3 ubiquitin ligase domain and domains involved in intra- and inter-molecular interactions (Figure 4A). The E3 ligase activity and protein interactions intertwine, which enables the ICP0 versatility when targeting the various anti-viral factors in host cells (26). To characterize the involvement of known ICP0 functions in MST1/2 cleavage and apoptosis, we infected HEp-2 cells with recombinant viruses containing the mCherry-tagged wild type or mutant ICP0 at different MOIs. Similar to that of the prototype HSV-1(F) shown in Figure 1, the recombinant virus containing the wild type ICP0 cDNA (RHG101) caused minimal cleavage of MST1/2 or PARP1 at 1 pfu/cell (Figure 4B, lane 2), indicating that the mCherry tag does not affect the mitigation of MST1/2 cleavage or apoptosis by ICP0. Intriguingly, the mutant virus with mutations in the E3 ubiquitin ligase of ICP0 (RHG120) triggered MST1/2 cleavage and PARP1 cleavage at 1 pfu/cell similar to that of the ΔICP0 virus infection (Figure 4B, lanes 3 vs. 8). Moreover, mutations in the SUMO-interaction motif (SIM) located at residues 362-364, which affect the binding of ICP0 to its SUMOylated substrates (RHG130) (32, 33), also induced MST1/2 cleavage and apoptosis but to a lesser extent than that of RHG120 (Figure 4B, lanes 3 vs. 5). Infection with a recombinant virus containing an N-terminal truncation in ICP0 (RHG105) had minimal production of MST1/2-NTs or PARP1 p89, whereas infection by the virus containing a C-terminal truncation in ICP0 (RHG103) generated substantial cleavage of both MST1/2 and PARP1 (Figure 4B, lanes 9 and 11), suggesting the different interaction partners linked to N- or C-terminus of ICP0 (26) have distinct roles in regulating the cleavage and apoptosis process. Moreover, the MST1/2 cleavage and apoptotic effects were overcome at 5 pfu/cell infection, similar to the dosage effects observed in the ΔICP0 infection (Figure 1B).

**Figure 4.**
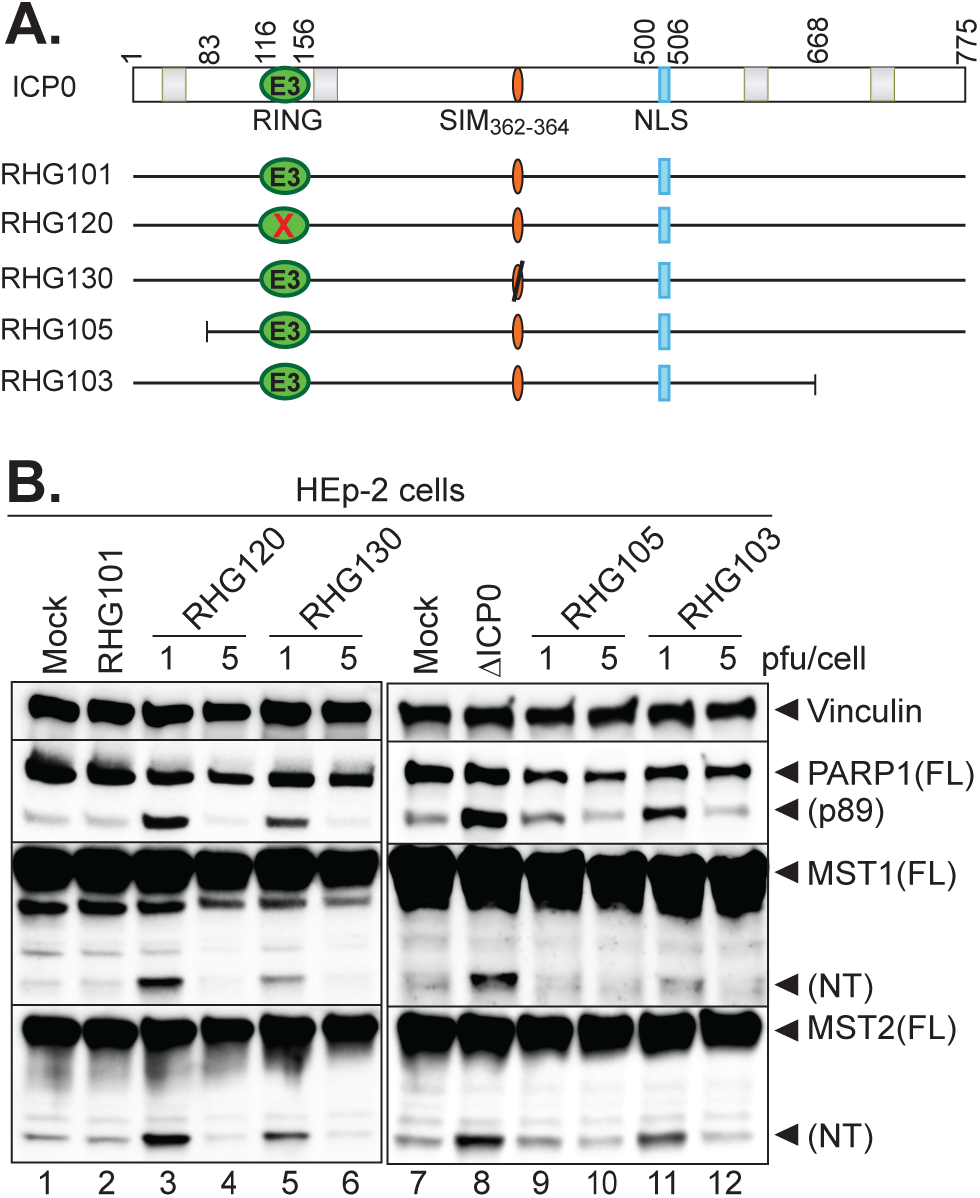
Known ICP0 functions intertwine with the Hippo reprogramming process. A) Schematic diagrams of ICP0 domains and mutations introduced into ICP0 in recombinant viruses RHG101, RHG120, RHG130, RHG105 and RHG103. Green oval with “E3”: RING-type E3 ubiquitin ligase domain; orange oval: SIM located at residues 362-364; blue rectangle: nuclear localization sequence; Red “X” in green oval: inactivated RING; black line cross orange oval: inactivated SIM; light grey boxes from left to right: binding sites for RNF8, cyclin D3, USP7, and CoREST as examples of ICP0 protein interactions. B) HEp-2 cells were infected by RHG101 or DICP0 virus at 1 pfu/cell, or by the indicated ICP0 mutant viruses at 1 or 5 pfu/cell for 24 hours before subjected to immunoblotting as described above.

### ICP0 executes disparate regulations on the cleavage of MST1 and MST2

The comparative experiments using U2OS and HEp-2 cells uncovered that ICP0 and US3 target different checkpoints for triggering MST1/2 cleavage and apoptosis (Figure 3). Since the pan-caspase inhibitor suppresses conversion of MST1-FL into MST1-NT (Figure 2E) and MST1/2 are direct substrates of caspase-1 and caspase-3 (17, 34), we used chemical inhibitors specific for caspase-1 or −3 to treat HEp-2 cells infected with different mutants. As wild type HSV-1 (F) possesses intact ICP0 and US3 to keep MST1/2 cleavage minimal, treatment with caspase-1 or caspase-3 inhibitor in HSV-1(F) infection had little effects (Figure 5). The caspase-3 inhibitor, Z-DEVD-FMK, significantly blocked the cleavages of PARP1 and MST1/2 in ΔUS3 virus infected cells (Figure 5, C, E, G), confirming that the activity of US3 in blocking intrinsic apoptosis and the correlation between MST1/2 cleavage and host apoptosis. Unexpectedly, an inhibitor specific to caspase-1, VX765, also significantly reduced the levels of cleaved PARP1 and MST1/2 in ΔUS3-infected cells, suggesting the involvement of this inflammatory caspase in the apoptotic pathway countered by US3 (Figure 5, B, D, F).

**Figure 5.**
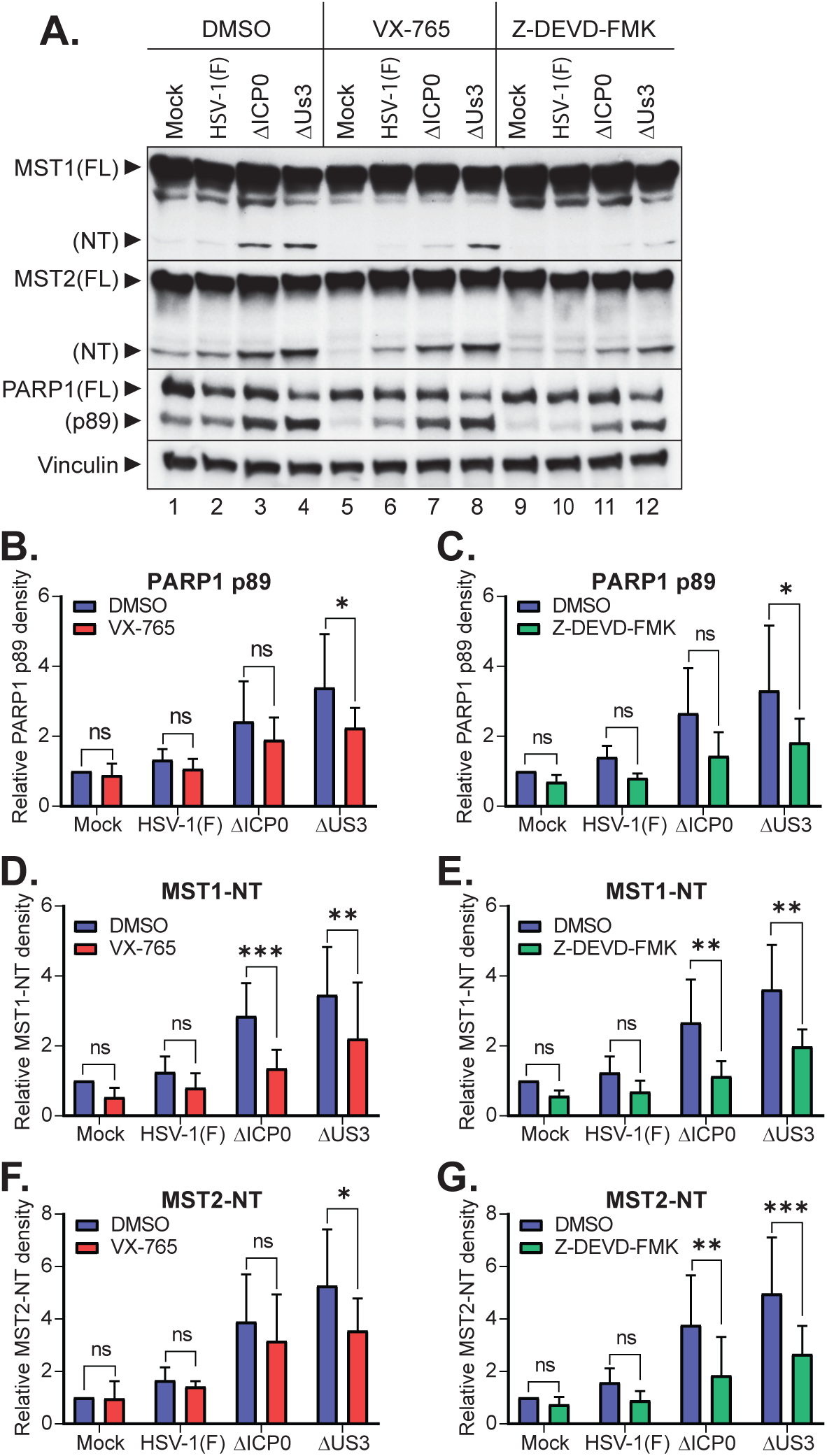
Differential inhibition on the ICP0- or US3-regulated MST1/2 and PARP1 cleavages by specific caspase inhibitors. HEp-2 cells were mock infected or infected with HSV-1(F), ΔICP0, or ΔUS3 virus at 1 pfu/cell for 24 hours in the presence of DMSO, caspase 1 inhibitor, VX-765, or caspase 3 inhibitor, Z-DEVD-FMK. A) Total cell lysates were immunoblotted for the indicated proteins. B-H) Band densities of PARP1 p89, and MST1/2-NT in VX-765 treatment (B, D, F) or Z-DEVD-FMK treatment (C, E, G) were normalized to vinculin. Fold changes compared to the Mock-DMSO sample were plotted in GraphPad. Data from 4 repeats were analyzed by two-way ANOVA. *: P<0.05; **: P<0.01; ***: P<0.001.

In an agreement with the role of ICP0 in counteracting MST1/2 cleavage and apoptosis, the caspase-3 inhibitor significantly reduced MST1/2-NT production (Figure 5, E, G) in HEp-2 cells infected with the ΔICP0 virus. Interestingly, the caspase-1 inhibitor significantly suppressed the production of MST1-NT while only marginally reduced MST2-NT (Figure 5, D, F), indicating that caspase-1 may have a preference toward MST1 over MST2 in the absence of ICP0. Although the levels of PARP1 p89 showed a decreasing trend upon the caspase-1 or caspase-3 treatment in ΔICP0-infected cells, the reduction was not statistically significant (Figure 5, B, C). Together, these results uncovered that caspase-1 and caspase-3 are both involved in cleaving MST1 and MST2 in HSV-1 infected cells and that viral proteins ICP0 and US3 differentially interfere with these caspases to prevent MST1/2 cleavage and apoptosis.

### Deletion of MST1/2 disrupts ΔICP0 infection at two different phases

To further define the role of MST1/2 in HSV-1 infection, we used CRISPR/Cas9 technology to knock out MST1/2 in HEp-2 cells. Human MST1 and MST2 share ∼75% sequence identity and many overlapped functions (2, 3). We isolated clones that had below detectable MST1 and MST2 proteins by immunoblotting, suggesting that these cell clones are MST1/2 double knockouts (MST1/2 DKO) (Figure 6A). The expression of other Hippo components, such as MOB1 and LATS1, were unchanged in the MST1/2-DKO clones, indicating that the knockout effect is specific for MST1/2. Moreover, while total MOB1 protein levels were unchanged, phosphorylation of MOB1 (pT35 MOB1) was minimal, further confirming the loss of MST1/2 since MOB1 is a direct substrate of MST1/2. Notably, MST1/2 DKO cells infected by HSV-1(F) had reduced PARP1 cleavage compared to the control HEp-2 cells transfected with a Cas9 plasmid without gRNA (Cas9) (Figure 6B and S1), indicating a critical role of MST1/2 in promoting the apoptotic responses during HSV-1 infection.

**Figure 6.**
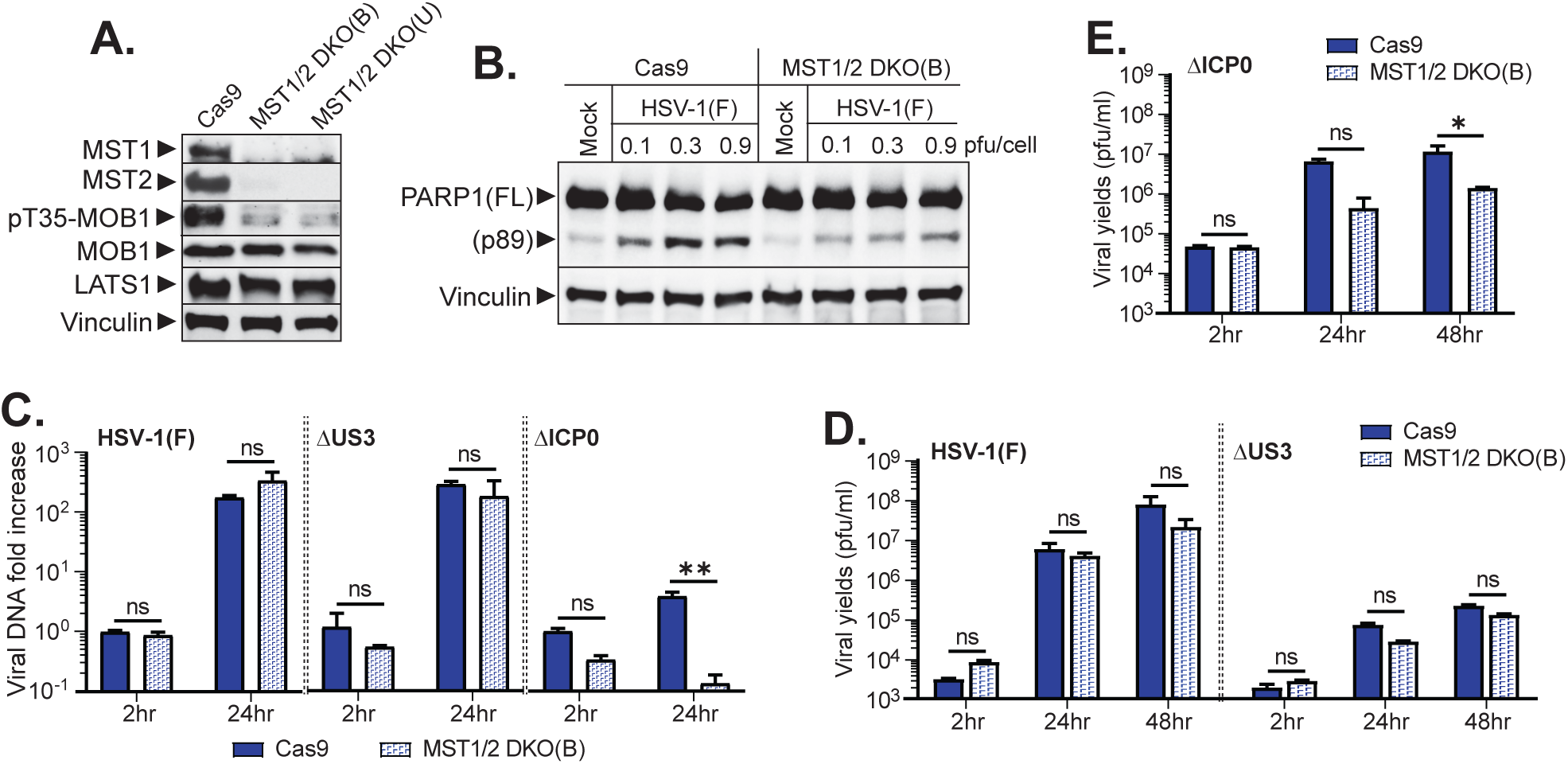
Double knockout of MST1/2 suppresses. ΔICP0 infection but shows marginal effects on that of HSV-1(F) or ΔUS3. A) One clone selected from the transfection of Cas9 only (Cas9) and two clones selected from the transfection of Cas9 plus MST1/2 gRNAs, MST1/2 DKO(B) and MST1/2 DKO(U), were grown, lysed and immunoblotted for MST1/2, MOB1, and LATS1 expression, and MOB1 phosphorylation.B) The Cas9 and MST1/2 DKO(B) cells were mock infected or infected by HSV-1(F) at 0.1, 0.3 and 0.9 pfu/cell for 24 hours and then immunoblotted for the indicated proteins.C) The Cas9 and MST1/2 DKO(B) cells were infected by HSV-1(F), ΔUS3 or ΔICP0 mutant at 0.3 pfu/cell. Total DNA extracted at 2 and 24 hours were subjected to comparative qPCR to measure viral DNA fold increase relative to the Cas9-2 hpi sample. D and E) The Cas9 and MST1/2 DKO(B) cells were infected by HSV-1(F) or ΔUS3 virus at 0.1 pfu/cell (D) or by ΔICP0 virus at 1 pfu/cell (E). At 2, 24 and 48 hpi, infected cells were lysed by sonication to measure the viral titer. Four repeats (C) or three repeats (D, E) were analyzed by two-way ANOVA (C, E) or three-way ANOVA (D). *: P<0.05; **: P<0.01.

Using the MST1/2 DKO(B) line, we tested the effects of deleting MST1/2 on HSV-1 replication. As expected, HSV-1(F) and the ΔUS3 virus robustly replicated with around 100-fold viral DNA increase in the Cas9 control cells, whereas the ΔICP0 virus replicated poorly in these cells with a mere 3.9-fold of DNA increase at 0.3 pfu/cell at 24 hpi (Figure 6C, Cas9). Surprisingly, the DNA fold increase of HSV-1(F) or ΔUS3 virus was not significantly changed in the MST1/2 DKO(B) cells, and the viral DNA abundance of ΔICP0 virus was reduced instead (Figure 6C, MST1/2 DKO(B)). In line with the results of DNA replication, the infectious virion production of HSV-1(F) or ΔUS3 virus was not significantly changed between Cas9 and MST1/2 DKO(B) cells, although the overall production of ΔUS3 virus was about 1.5-2 log less than that of HSV-1(F) using the 0.1 pfu/cell input (Figure 6D). Furthermore, the virion production of ΔICP0 virus at 1 pfu/cell input showed a ∼10-fold reduction in MST1/2 DKO(B) cells compared to that of Cas9 cells, indicating a requirement of MST1/2 in the ΔICP0 infection (Fig. 6E).

To validate the importance of MST1/2 in ΔICP0 infection, we compared the ΔICP0 DNA replication in another MST1/2 double-knockout clone, MST1/2 DKO(U). To ensure that equivalent initial MOI is applied to multiple cell lines, we evaluated 0 hpi sample as shown in the experimental timeline (Figure 7A). Surprisingly, we observed an drastic reduction in the viral DNA level from 0 hpi to 2 hpi in the two MST1/2 DKO cell lines compared to Cas9 at 1 pfu/cell but not at 5 pfu/cell (Figure 7B and 7C), indicating a dosage-dependent viral DNA clearance for the ΔICP0 virus immediately after the viral entry in the absence of MST1/2. Furthermore, the defective ΔICP0 DNA replication at 24 hpi in MST1/2 DKO cells persisted at both 1 and 5 pfu/cell (Figure 7C and 7D), suggesting a dosage-independent defect in the DNA replication step of ΔICP0 infection in the absence of MST1/2.

**Figure 7.**
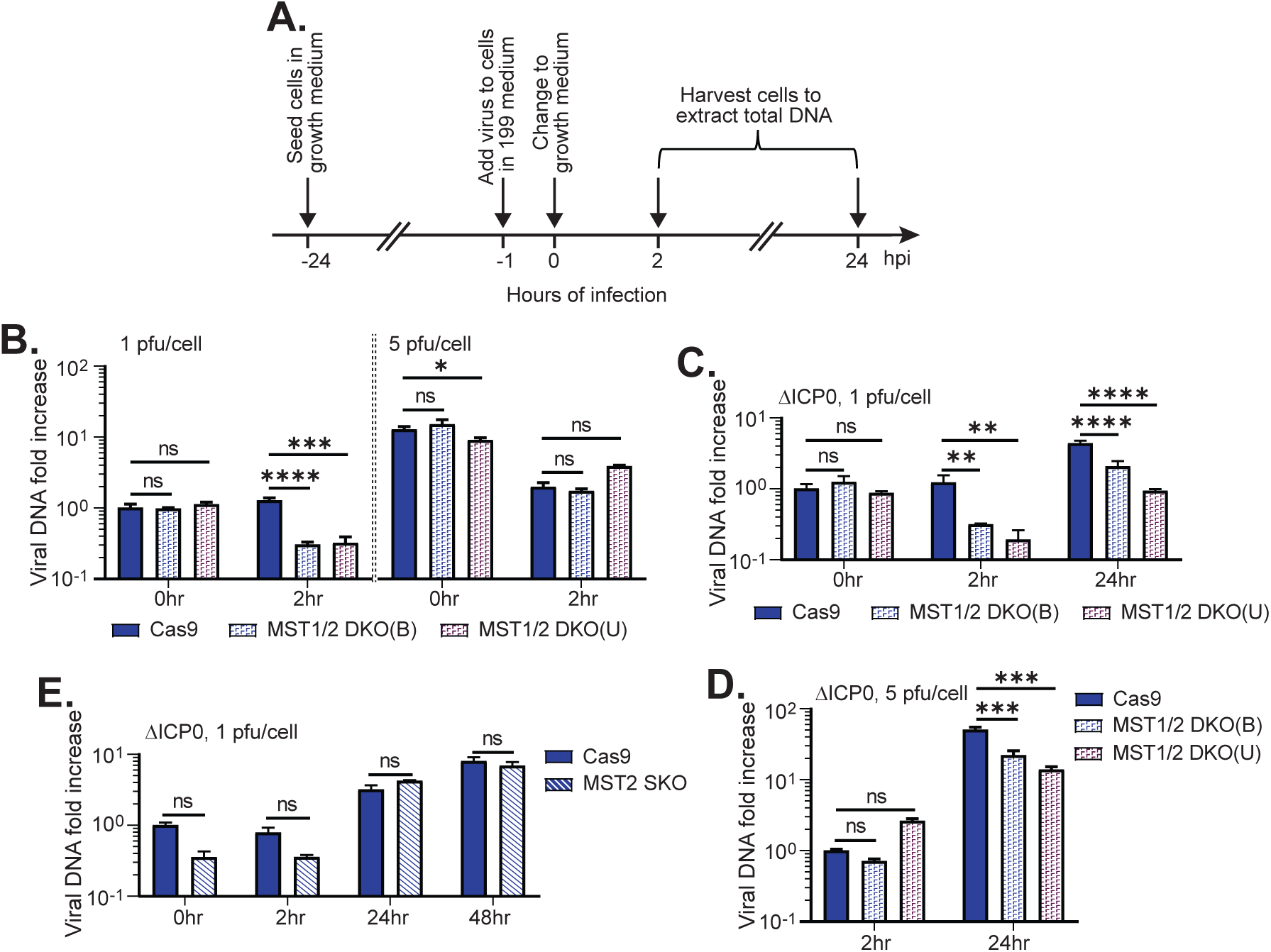
Double knockout of MST1/2 suppresses. ΔICP0 infection in dosage-dependent and dosage-independent manners. A) Detailed experimental timeline. B-D) The Cas9, MST1/2 DKO(B) and MST1/2 DKO(U) cells were infected by ΔICP0 virus at 1 or 5 pfu/cell. Total DNA extracted at 0, 2 and 24 hours as shown in (A) were subjected to comparative qPCR to measure the viral DNA fold increase relative to the Cas9-0 hpi sample (B, C), or relative to the Cas9-2 hpi sample (D). E) The Cas9 and MST2 SKO cells were infected by ΔICP0 virus at 1 pfu/cell. Total DNA extracted at 0, 2 and 24 hours were subjected to comparative qPCR to measure the viral DNA fold increase relative to the Cas9-0 hpi sample. Three repeats were analyzed by two-way ANOVA. *: P<0.05; **: P<0.01; ***: P<0.001; ****: P<0.0001.

MST1 and MST2 are highly homologous and functionally redundant in controlling cell growth and tumorigenesis (35, 36). To further characterize the importance of MST1/2 in ΔICP0 infection, we constructed single knockout (SKO) cell lines by transfecting the individual MST1/2 gRNAs. However, transfection of MST1 gRNA alone drastically reduced both MST1 and MST2 expression (Figure S2, lanes 9-12). Therefore, we only conducted analysis using the MST2 single knockout cells. We again noticed the lowered PARP1 cleavage in HSV-1(F) infection (Figure S2, lane 2 vs 6), suggesting that MST2 by itself mediates an apoptotic reaction in HSV-1(F) infection that cannot be complemented by the MST1 activities. Furthermore, ΔICP0 DNA level remained consistent from 0 hpi to 2 hpi in both Cas9 and MST2 SKO cells, whereas the ΔICP0 DNA replication in MST2 SKO cells was also executed as effective as that in Cas9 cells (Figure 7E). These results suggest that the presence of MST1 compensates for the loss of MST2 in stabilizing ΔICP0 DNA at the immediate early stage and facilitating the ΔICP0 DNA replication in a later stage.

## Discussion

The Hippo pathway is a conserved signaling pathway important for many critical processes including cell growth, organ development and cancer suppression. In light of the findings regarding recurrent HSV infections in MST1-deficient patients (4), this study investigated the unexplored role of MST1/2 in human epithelial cells challenged with HSV-1. Beyond our previous study reporting MST1/2 cleavage in immune cells upon detection of bacterial pathogens and chemical stimuli (16, 17), the present study discovered that MST1/2 cleavage and the apoptosis hallmark, PARP1 cleavage, occur in HEp-2 cells, a non-immune cell type, in response to HSV-1 (Figure 1). Moreover, ectopic expression of MST1 cleavage products prior to infection significantly constrained the infection in the aspects of viral protein expression, DNA replication, and virion production (Figure 2). This is the first study to demonstrate the proteolytic cleavage of MST1/2 in association with viral infection in epithelial cells, revealing the generality of Hippo reprogramming in both immune and non-immune cells as an autonomous host response against the intracellular bacterial and viral pathogens. The mild cleavage detected in HSV-1(F) infection is substantially enhanced when either ICP0 or US3 gene is deleted and the amount of cleavage is regulated in a dosage dependent manner, revealing that the host defense triggered by MST1/2 cleavage can be saturated by increasing certain viral functions such as the immediate early protein ICP0 and the late protein US3 (Figure 8).

**Figure 8.**
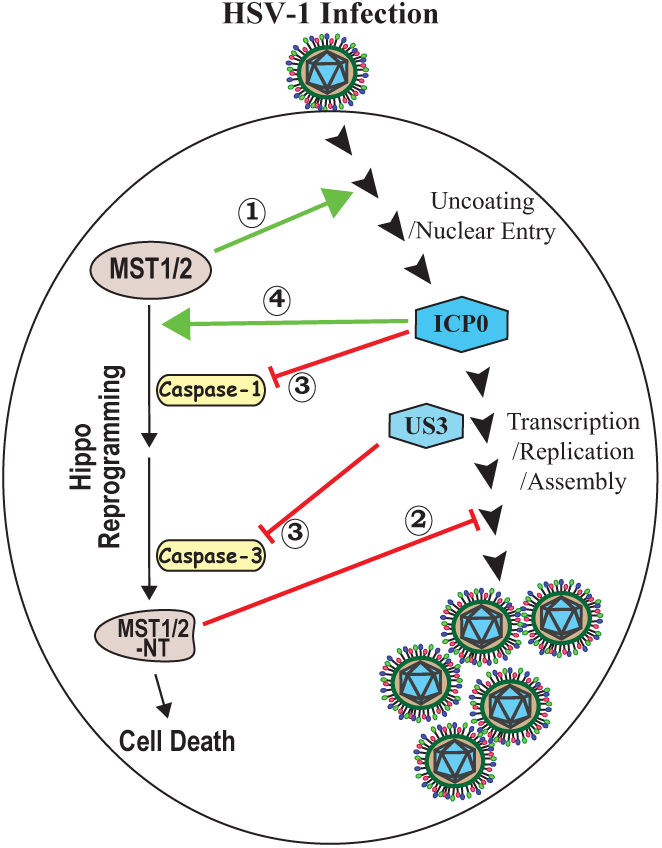
Multiple levels of interactions between HSV-1 and the full-length or cleaved MST1/2. The major HSV-1 infection steps are illustrated on the right side, with ICP0 and US3 proteins represented by hexagons.: Full-length MST1/2 facilitate the early viral DNA intake;: Cleaved MST1/2-NT variants suppress viral replication;: Caspase-1 acts upstream of Caspase-3, whereas ICP0 and US3 target caspase-1 caspase-3, respectively;: ICP0 expression by itself promotes the host Hippo reprogramming.

Interestingly, deletion of both MST1 and MST2 genes did not improve HSV-1 replication. Instead, we found suppressions on the ΔICP0 virus at multiple infection phases in MST1/2 DKO cells and showed only marginal effects on HSV-1(F) or ΔUS3 infection (Figure 6). This is not unprecedented because several host pathways have both pro-viral and anti-viral roles in HSV-1 infection, especially those essential for cell cycle regulation (37, 38). The results from MST1/2-NT overexpression and MST1/2 double deletion might seem opposite but, in fact, suggest that the full-length MST1/2 and the canonical Hippo signaling pathway may facilitate HSV-1 infection, the lack of which can be overcome in the HSV-1(F) and ΔUS3 infection but not in infection without ICP0 (Figure 7). As a counter measurement to avoid being exploited by the virus, the host cells activate the Hippo reprogramming process which irreversibly cleaves the full-length proteins into the pro-apoptotic MST1/2-NT fragments (Figure 8). Further examination of other Hippo components, identification of the phosphorylation substrates of MST1/2, and characterization of the regulations on Hippo kinases, in the context of MST1/2 cleavage will help to elucidate the relations between the canonical Hippo pathway and the noncanonical Hippo reprogramming during HSV-1 infection.

Our inquiries of HSV-Hippo interactions not only show that full-length MST1/2 and their cleaved variants affect HSV-1 infection on different levels but also reveal that HSV-1 imposes multifaceted regulations to manipulate the MST1/2 cleavage and host apoptosis. The first level of complexity in the HSV-Hippo interactions is the deployment of multiple viral proteins, of which we identified ICP0 and US3 so far, to counter MST1/2 cleavage and apoptosis during HSV-1 infection. It is worth noting that various HSV-1 gene products have been reported to block apoptosis, including US3 (18, 19), gD (39, 40), gJ (39, 41), and LAT (42, 43). Potentially, additional viral proteins or viral transcripts other than ICP0 and US3 may intertwine in the MST1/2 cleavage and apoptosis triggered by HSV-1 infection.

The second layer of complexity in HSV-1 counteraction is reflected in the distinctive mechanisms used by ICP0 and US3 in their regulations on the Hippo reprogramming. The results show that US3, a well characterized apoptosis blocker, also prevents MST1/2 cleavage. Unlike US3, ICP0 plays dual roles in MST1/2 cleavage and host apoptosis (Figure 3). In the infection context, ICP0 executes anti-apoptotic functions to block the accumulation of MST1/2-NT and PARP1 p89 in non-permissive HEp-2 cells, but not in the permissive U2OS cells, where certain restrictive factors targeted by ICP0 are defective. Our results that ΔICP0 infection enhanced MST1/2 cleavage and apoptosis in HEp-2 cells but not in U2OS cells link the Hippo programming process to the defective chromatin repression and STING/IFN response pathways in U2OS cells (30, 31).

The interesting part is that overexpression of ICP0 alone in both HEp-2 and U2OS cells induces MST1/2 cleavage and apoptosis (Figure 3A and 3B). The pro-apoptotic activity of ICP0 is likely the outcome of ICP0 targeting host intrinsic defenses. For example, the first substrates identified for the ICP0 E3 ubiquitin ligase are PML (promyelocytic leukemia) protein and Sp100 (speckled protein 100) (44), both of which are core components of the dynamic nuclear structures called ND10s or PML-NBs (45, 46). ND10s are crucial in genome integrity, epigenetic regulation, and cell apoptosis (47–49). During HSV-1 infection, the ICP0 E3 mediated PML and Sp100 degradation disperses ND10s to promote viral gene expression and DNA replication (50, 51). ICP0 expressed via transfection attacks ND10s, causing disturbance to the ND10 structure without the infection. Although the mechanism remains unexplored, it is plausible to postulate that the loss of ND10 integrity acts as an apoptosis inducer and contributes to the pro-apoptotic activity of ICP0 (Figure 8). Whether the activation of MST1/2 cleavage and apoptosis benefits HSV-1 at the immediate early stage of infection or how nuclear-cytoplasmic communication enables Hippo components in the cytoplasm sense the ND10 disruption in the nucleus are questions critical to understanding the HSV-host interactions.

Dual actions of ICP0 in regulating host pathways have been previously reported, where results show that ICP0 counteracts a self-guarded anti-viral defense executed by MORC3 (microrchidia 3) with two different functions (52). MORC3 is a nuclear protein localized at ND10 that influences the recruitment of ATRX/Daxx complex to HSV-1 genome and also a negative regulator of IFN production (52, 53). Upon infection, ND10 components converge with the incoming DNA to force the naked DNA into viral chromatin and repress viral transcription. The proteasomal degradation of MORC3 mediated by ICP0 E3 releases the chromatin repression but also unleashes IFN synthesis to place additional restrictions on viral replication, which is again negated by ICP0 functions, along with other viral proteins (52, 54, 55). The dual roles of ICP0 in the apoptosis triggered by MST1/2 cleavage may be analogous to the ICP0-MORC3 relation, through which an early ICP0 action may activate certain caspases to trigger Hippo reprogramming and the apoptotic responses resulted from MST1/2 cleavage further requires ICP0 and US3 functions to overcome in late infection.

The third layer of complexity in ICP0-Hippo interactions is the involvement of multiple caspases in MST1/2 cleavage and the differential regulations on MST1 and MST2 cleavage in the ΔICP0 infection. The caspase inhibition experiment showed that specific inhibitors for caspase-1 or caspase-3 significantly reduced MST1/2 cleavage or PARP1 cleavage, but in a context-dependent manner during the ΔICP0 or ΔUS3 infection (Figure 5). PARP1 cleavage as one of the outcomes of caspase-1 activation by the inflammasomes in macrophages has been reported (17, 56, 57). Our results show that both caspase-1 and caspase-3 inhibitors diminish MST1/2 and PARP1 cleavage in ΔUS3 infected cells, while neither blocked PARP1 cleavage in ΔICP0 infection. These interesting observations show that caspase-1 triggered PARP1 cleavage also exists in non-immune cells and that caspase-1 may act upstream of the apoptotic reactions blocked by US3 (Figure 8). The caspase-1 inhibitor reduces MST1 cleavage but had no effects on MST2 cleavage in ΔICP0 infection (Figure 5D-G), which aligns with the notion that ICP0 has multifaceted roles in the interaction with the Hippo pathway. We have previously reported that ICP0 has the ability to differentiate host proteins that share high similarity, such as PML isoforms that are only different in their C-terminus, and target their degradation by distinctive biochemical mechanisms (32, 58). Whether this type of host protein differentiation by ICP0 is coordinated with MST1/2 reprogramming will be an important project to continue in understanding the anti-viral properties of Hippo kinases.

## Acknowledgements

We thank the anonymous reviewers for their constructive feedback. The research was supported by the Wayne State University Research Grant and the Startup Funds awarded to Pei-Chung Lee, and Wayne State University Bridge Funding I awarded to Haidong Gu.

## Materials and Methods

### Cells and Viruses

Human epithelial HEp-2 cells and African green monkey kidney Vero cells were cultured in Dulbecco’s modified Eagle medium (DMEM) (Invitrogen) supplemented with 10% fetal bovine serum (FBS). Human osteosarcoma cells U2OS were grown in McCoy’s 5A medium supplemented with 10% FBS. The prototype HSV-1(F) strain and its derived R7041(ΔUS3 virus) (59) were provided by Dr. Bernard Roizman at University of Chicago and amplified on Vero cells. Recombinant viruses RHG101, RHG120, RHG130, RHG105, RHG103 and R8501 have been described elsewhere (60–62).

### Restoration of thymidine kinase (TK) gene in the ICP0-null virus R8501

ICP0-null virus R8501 (62) is constructed via a bacterial artificial chromosome (BAC) inserted in the TK gene of HSV-1(F). To restore TK in R8501, we established a U2OS-TK^-^ cell line. Primers 5’-GATCGGATCTGCCCCCGGGTCTTGCG-3’ and 5’-AAAACGCAAGACCCGGGGGCAGATCC-3’ were annealed and then inserted into the BamHI and BsmBI digested pCas-Guide (OriGene). The resulted pCas-TK1 gRNA plasmid was transfected into U2OS cells and selected for U2OS-TK^-^ cells in McCoy’s 5A medium supplemented with 10% FBS and 100 μg/ml of 5-Bromodeoxyuridine (Sigma, B5002). Stable U2OS-TK^-^ cells were then transfected with plasmid pRB4867 (63) for 24 hours before being infected with R8501 at 0.1 pfu/cell. The TK^+^ progeny, ΔICP0 virus (RHG501), was plaque purified and amplified in McCoy’s 5A medium supplemented with 10% FBS, 1x Hypoxanthine-thymidine (Sigma, H0137) and 20 μg/ml of aminopterin (Sigma, A3411).

### Construction of MST1 and MST2 knockout cell lines

Guide RNA corresponding to either human MST1 (5’-TGGATCGTTATGGAGTACTG-3’) or MST2 (5’-TGGATTGTTATGGAGTACTG-3’) were cloned into the pSpCas9(BB)-2A-Puro (PX459) V2.0 plasmid (Addgene #62988). HEp-2 cells were seeded one day before transfection in a 6-well plate at 5×10^5^ per well. 2µg of DNA was transfected into wells using Lipofectamine 2000 diluted in Opti-MEM (ThermoFisher). 24 hours post-transfection, cells were washed and incubated with media containing 1µg/mL puromycin. The puromycin selection occurred for 3-4 days, following which, cells were allowed to recover for 24 hours in complete DMEM without puromycin. Finally, attached cells were washed from the wells, diluted to 2×10^1^, and seeded into a 96-well plate to allow single colonies to grow. Knockout clones from these single colonies were validated with western blotting.

### Western Blotting

Cells transfected and/or infected were harvested, washed with phosphate-buffered saline (PBS) and lysed in a radioimmunoprecipitation assay (RIPA) buffer (50 mM Tris, pH7.4, 150 mM NaCl, 1 mM EDTA, 0.1% SDS, 1% NP40, 0.25% Sodium Deoxycholate) by sonication. Total cell lysates were denatured with Laemmli buffer, separated by SDS-polyacrylamide gel electrophoresis and then transferred to a polyvinylidene difluoride (PVDF) membrane (Thermo Scientific, Rockford, IL, USA) or nitrocellulose membrane (Bio-Rad). The membrane was blocked with 1× Tris-buffered saline–Tween (TBST) (20 mM Tris, pH7.5, 150 mM NaCl, 0.5% Tween 20) containing 5% nonfat dry milk, and then probed with a primary antibody, rinsed with TBST for three times to react with the horseradish peroxidase-conjugated goat anti-mouse or goat anti-rabbit secondary antibody (Sigma, St. Louis, MO, USA). The membrane was finally rinsed with TBST and visualized by the ECL detection reagent (Thermo Scientific) and imaged with Bio-Rad chemiluminescent imager (Chemi-Doc) or iBright (Invitrogen). Band density from the Chemi-Doc exposure was quantitated by the Bio-Rad Image Lab software, and that from iBright exposure was quantitated by Image J.

### Comparative qPCR to measure viral DNA replication

Cells infected with a wild-type or recombinant HSV-1 at indicated MOI was harvested at indicated hpi, washed with PBS and digested by 2 mg/ml proteinase K (Roche) at 55°C overnight before treating with 50 μg/ml RNase A. Total DNA was isolated from the digested lysates by phenol-chloroform extraction and subjected to quantitative PCR (qPCR). For viral DNA quantification, the ICP27 gene was targeted using primers 5′-GCAGCTAGCATGGCGACTGACATTGATATG-3′ and 5′-GCAGAATTCCTAAAACAGGGAGTTGCAATA-3′ in a Thermo Fischer Quant Studio 3 thermocycler. Universal 18S internal primer mix from Invitrogen Life Technologies was used as internal reference. Δ*C_T_* was normalized to the viral input detected at 0 or 2 hpi to calculate viral DNA fold increase 2^(-ΔΔ^*^CT^*^)^.

### Viral titration

To measure viral growth, HEp-2 cells were infected with the wild type or mutant HSV-1 at 0.1 pfu/cell and harvested at timepoints. Cells resuspend in milk were frozen and thawed, followed by sonication to release the virion. Viral growth of HSV-1(F) or ΔUS3 virus was titrated on Vero cells, and that of ΔICP0 virus was titrated on U2OS cells in the presence of human gammaglobulin (Sigma Aldrich, USA). Viral plaques were fixed with methanol and stained with 1X Giemsa Staining. Plaques from triplicated experiments were counted and plotted with GraphPad.

### Plasmids

Human full-length and N-terminal (1–326) cleaved fragments for both MST1 and MST2 were cloned into a mammalian expression plasmid containing a CMV promoter (backbone before removal of Flag tag: Addgene #1965). Plasmids expressing ICP0 (pAc-CMV-ICP0) or US3 (pAc-CMV-US3), and pRB4867 were generous gifts from Bernard Roizman at University of Chicago (63–65).

### Antibodies

Antibodies for vinculin and actin were purchased from Santa Cruz Biotechnology (sc-73614; sc-81178). Antibody for MST2 was purchased from Abcam (ab52641). Antibodies for PARP1, MST1, MOB1, pT35-MOB1, and LATS1 were purchased from Cell Signaling (9542S; 14946S; 13730S; 8699S; 3477S). Antibodies for IC27, ICP8, and gC were purchases from Virusys. Antibody for US3 was a generous gift from Dr. Bernard Roizman at University of Chicago (66). Production of rabbit polyclonal antiserum against ICP0 has been described elsewhere (67).

### Reagents

Staurosporine was purchased from Cayman Chemical (#81590). Raptinal was purchased from Adipogen (AG-CR1-2902). Z-VAD-FMK and Z-DEVD-FMK were purchased from R&D Systems. VX765 was purchased from InvivoGen (inh-vx765i-1)

